# Near completely reversing the γ- to β-globin switch by enhancer release, retargeting and reinforcing

**DOI:** 10.64898/2026.01.30.702713

**Authors:** Nanyu Wang, Ke Yang, Xuemin Xie, Shenshen Cui, Xiaoya Pei, Xiang Zhao, Delong Hao, Yuyan Jia, Gaohui Yang, Rongrong Liu, Ping Chen, Wenji Dong, Yue Huang, Xiang Lv, Zhuqin Zhang, Depei Liu

## Abstract

The γ- to β-globin switch is intricately regulated during human ontogeny, and this process is manipulated for therapeutic approaches to treat β-hemoglobinopathies by activating γ-globin expression. Several genetic strategies to reactivate HbF have partially reversed the γ- to β-globin switch and ameliorated the clinical symptoms of β-hemoglobinopathies. However, whether the γ- to β-globin switch can be completely reversed remains unknown. Completely reversing the γ- to β-globin switch requires a thorough redirection of the locus control region (LCR) from interacting with the β-globin gene (*HBB*) to interacting with the γ-globin gene (*HBG*). Here, we found that disrupting the KLF1-mediated *HBB*-LCR interaction by mutating the CACCC motif in *HBB* leads to the release of the LCR and its retargeting to other β-like globin genes. Moreover, simultaneously disrupting the KLF1-mediated *HBB*-LCR interaction and the epigenetic repression of *HBG* by combined editing of the CACCC motif in *HBB* and the TGACCA motif in *HBG* reinforces the *HBG*-LCR interaction, resulting in almost exclusive γ-globin expression while nearly absent β-globin expression, achieving near complete reversal of the γ- to β-globin switch. This finding demonstrates the comprehensive regulation of the γ- to β-globin switch by gene competition and gene silencing mechanisms. This finding also suggests that silenced genes can be fully activated through the redirection of enhancer-promoter contacts and that the specificity of enhancer-promoter contact within chromosomal domains is achieved through the transcription factor clusters binding to enhancers and promoters. Combined editing of the CACCC&TGACCA motifs also offer a more optimal therapeutic strategy for β-hemoglobinopathies.

## Introduction

During human development, the major form of hemoglobin in red blood cells changes from fetal hemoglobin (HbF, α2γ2) to adult hemoglobin (HbA, α2β2) with the migration of hematopoietic sites^1^. This transcriptional shift from the γ-globin gene (*HBG1/HBG2*, referred to as *HBG*) to the β-globin gene (*HBB*) is commonly referred to as the γ-to β-globin switch. Mutations in *HBB* cause β-hemoglobinopathies, including β-thalassemia and sickle cell disease (SCD), which are the most common monogenic diseases and impose significant burdens on public health^2,3^. Reactivating HbF expression in adults can alleviate the clinical symptoms of β-hemoglobinopathies and has emerged as a potential therapeutic approach^4,5^. Currently, several genetic approaches to reactivate HbF expression, including relieving the epigenetic silencing of *HBG*^*6-13*^, forcing chromatin looping to *HBG*^14^ and disrupting the *HBB* promoter^15^, have achieved partial reversal of the γ-to β-globin switch. Variations in disease severity among populations with hereditary persistence of fetal hemoglobin (HPFH) indicate that as HbF levels further increase, the clinical manifestations in patients with β-hemoglobinopathy become milder, necessitating further elevation of HbF levels^16,17^. However, whether the γ-to β-globin switch can be completely reversed remains unknown.

As a super-enhancer, the locus control region (LCR) increases globin gene expression by forming a chromatin loop, thereby facilitating its close physical proximity to globin loci^18-20^. The LCR shifts its proximity from *HBG* to *HBB* during the γ-to β-globin switch^19^. Hence, completely reversing the γ-to β-globin switch requires a thorough redirection of the LCR from interacting with *HBB* to interacting with *HBG*. Previous studies have forced the LCR to loop to *HBG*, effectively activating HbF expression, but didn’t achieve thorough redirection of the LCR^14^. Here, we show that disrupting the KLF1-mediated *HBB*-LCR interaction by disrupting the CACCC motif in *HBB* leads to the release of the LCR and its retargeting to other β-like globin genes. Moreover, simultaneously disrupting the epigenetic silencing of *HBG* and the KLF1-mediated *HBB*-LCR interaction by combined editing of the CACCC motif in *HBB* and the TGACCA motif in *HBG* reinforces the interaction between *HBG* and the LCR, achieving near complete reversal of the γ-to β-globin switch. This finding not only demonstrates the comprehensive regulation of the γ-to β-globin switch by gene competition and gene silencing mechanisms and broadens our understanding of eukaryotic gene regulatory mechanism, but also offers a more optimal therapeutic strategy for β-hemoglobinopathies.

## Results

### LCR release and retargeting by disrupting the KLF1-mediated *HBB*-LCR interaction

The LCR interacts with *HBB* in adults, so redirecting it from its proximity to *HBB* towards *HBG* requires first releasing it from its interaction with *HBB*. The functional loss of a preferred promoter releasing its partner enhancer to loop to and activate alternative promoters^21^, as well as the disruption of *HBB* promoter activity activating *HBG* expression^15^, together indicate that the release of the LCR from its interaction with *HBB* reactivates *HBG* expression. To identify the regulatory elements that can be disrupted to achieve the release of the LCR from its interaction with *HBB*, we performed a CRISPR-Cas9 screen in HUDEP-2 cells, which is an immortalized cell line that primarily expresses HbA^22^. A lentiviral library encoding 3,761 single-guide RNAs (sgRNAs) targeting downstream regions of *HBG* (hg38/GRCh38, chr11: 5204054-5248601) was transduced into Cas9-expressing HUDEP-2 cells, 24 of which were designed to target homologous regions of *HBB* and *HBD*. The cells were then sorted based on HbF expression, and sgRNA sequences in the genomic DNA were analyzed by next-generation sequencing (NGS) (Supplementary information, Fig. S1a). Among the sgRNAs enriched in HbF^high^ cells, the top seven targeted the homologous regions of *HBB* and *HBD* (containing the *HBB* promoter in their intervening region), followed by the proximal CACCC motif of KLF1 in the *HBB* promoter (Fig. 1a; Supplementary information, Fig. S1b). These findings are consistent with previous studies showing that KLF1 is a major transcription factor responsible for activating β-globin expression in adults, and is essential for the formation of an active chromatin hub in the β-globin gene cluster, as well as for the construction of a chromatin conformation that facilitates the interaction between *HBB* and the LCR^23-27^. KLF1 has two binding sites in *HBB*, the proximal CACCC motif and the distal CACCC motif^28,29^ (also enriched in the CRISPR screen (Fig. 1a)). Notably, a previous study of the disruption of *HBB* promoter activity activating *HBG* expression also found that disrupting these two CACCC motifs could increase *HBG* expression^15^. And mutations in these motifs increased HbF levels to 1.3-65% of total hemoglobin levels in individuals^30^ (Fig. 1b). We electroporated HUDEP-2 cells with ribonucleoprotein (RNP) complexes carrying either PM-1 and PM-2, which target the proximal CACCC motif in *HBB*, or DM, which target the distal CACCC motif in *HBB*. The editing of the CACCC motif increased the expression of *HBE, HBG* and *HBD*, suggesting that after the LCR was released from its interaction with *HBB* by disrupting the CACCC motif, it retargets not only to *HBG* but also to *HBE* and *HBD* (Fig. 1c; Supplementary information, Fig. S1c). The electroporation of PM-1, PM-2 and DM into Cas9-expressing HUDEP-2 cells also increased the proportion of F cells from 3% to 35%, 21%, and 15%, respectively (Supplementary information, Fig. S1d). To determine whether mutation of the CACCC motif in *HBB* causes the release of the LCR from interacting with *HBB* and its retargeting to interact with other β-like globin genes, we performed a chromosome conformation capture (3C) assay and found that editing the CACCC motif weakened the *HBB*-LCR interaction while enhanced the *HBG*-LCR interaction (Fig. 1d). Additionally, the regulation of target genes by enhancers can alter their transcriptional bursting frequency^31,32^. We observed that editing the CACCC motif decreased the transcriptional bursting frequency of *HBB*, while increased that of *HBG* in HUDEP-2 monoclonal cells (Fig. 1e; Supplementary information, Table S1). Previous study has shown that the binding of CTCF to promoters determines the specificity of enhancer-promoter interactions^21^, while CTCF binds only to the flanking regions on two sides of the β-globin cluster^33^, and the binding of CTCF within the β-globin cluster remained unchanged after the CACCC motif was edited (Supplementary information, Fig. S1e). We observed that editing the CACCC motif reduced chromatin accessibility, RNAPII binding and the active transcriptional epigenetic modification H3K4me3 and H3K9Ac on *HBB*, while slightly increased that on *HBG*, accompanied by changes in the binding of transcription factor complexes to the β-globin gene cluster^9,13,26,27,34^ (Fig. 1F; Supplementary information, Fig. S1e). These results suggest that the release of the LCR from *HBB* and its retargeting after CACCC motif mutation is mediated by changes in the cluster of transcription factors bound to the promoter.

**Fig 1.**
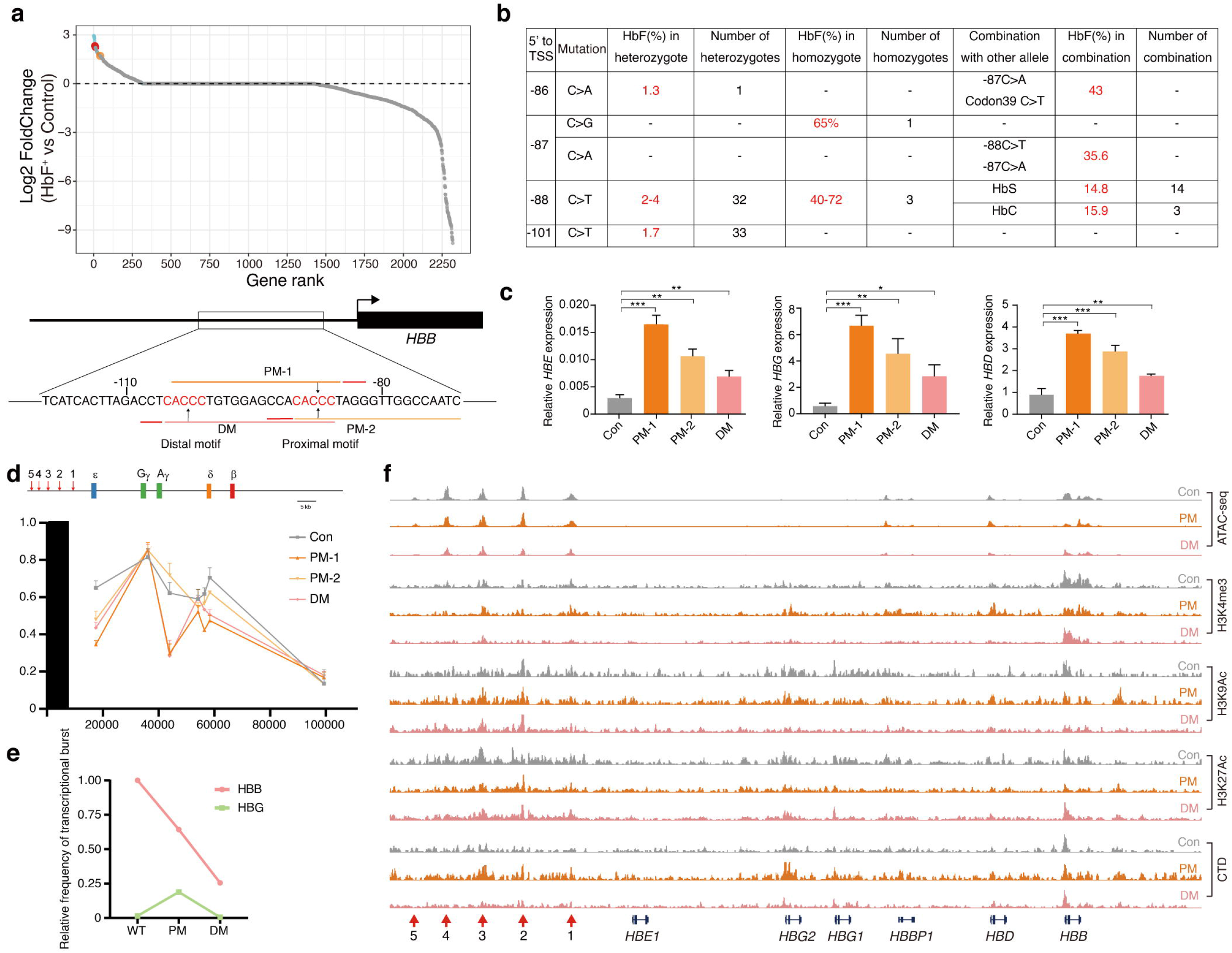
LCR release and retargeting by disrupting the CACCC motif in *HBB* promoter. **a** The results of the CRISPR–Cas9 screen showing enriched sgRNAs in HUDEP-2 cells expressing high levels of HbF. The y-axis shows the log_2_(fold-change (FC)) in sgRNAs. The sgRNAs targeting the homologous sequences of *HBB* and *HBD* (blue dots), the proximal CACCC motif (red dot) and the distal CACCC motif (orange dot) in *HBB* are highlighted. **b** HbF levels of individuals with mutations in the CACCC motif in the *HBB* promoter in the Globin Gene Server. **c** Cas9-expressing HUDEP-2 cells were transduced with sgRNAs targeting the CACCC motif in *HBB*. The average editing efficiencies of PM-1, PM-2 and DM were 79.1%, 67.4% and 45.4%, respectively. The charts show β-like globin gene expression relative to β-actin mRNA expression as measured by RT–qPCR (mean ± s.d., n = 3). Multiple comparisons were assessed with one-way ANOVA with Tukey’s MCT. \**P* < 0.05, \*\**P* < 0.01 and \*\*\**P* < 0.001. **d** Chromosome conformation capture analysis of control and CACCC motif edited HUDEP-2 cells (mean ± s.d., n = 3). **e** The relative frequencies of the transcriptional bursts of *HBB* and *HBG* in HUDEP-2 clones were tested by Chromium single cell sequencing. **f** ATAC-seq signals at the β-globin cluster were analyzed in control and CACCC motif edited HUDEP-2 cells, along with CUT&Tag enrichment for CTD, H3K4me3, H3K9Ac, and H3K27Ac.

### Near complete reversal of the γ-to β-globin switch by reinforcing the HBG-LCR interaction

After mutation of the CACCC motif in *HBB*, the LCR is released from its interaction with *HBB* and retargeting to other β-like globin genes, but the interaction between other β-like globin gene and the LCR is somewhat weak (Fig. 1d, e). In adults, *HBG* is silenced by many suppressors, especially BCL11A, which is one of the major HbF silencers, and recruits many epigenetic repressors such as the nucleosome remodeling and deacetylase (NuRD) complex, to silence γ-globin expression^9^. We hypothesized that releasing this suppression may reinforce the *HBG*-LCR interaction and further increase γ-globin expression. By disrupting the distant TGACCA motif of BCL11A, the epigenetic repression of *HBG* can be relieved^11,12^. Therefore, we first edited the TGACCA motif in *HBG* in HUDEP-2 cells (GM) and subsequently edited the CACCC motif in *HBB* in this edited cell population to obtain cells with simultaneous editing of both the CACCC motif in *HBB* and the TGACCA motif in *HBG* (PG or DG) (Fig. 2a). Compared with editing the TGACCA motif alone, editing both motifs simultaneously increased the expression of γ-globin by 1.9-fold (PG) and 1.8-fold (DG) (Fig. 2B). The level of HbF also increased from 8.6 mAU*min to 18.2 mAU*min and 16.8 mAU*min, respectively (Supplementary information, Fig. S2a). The proportion of F cells also consistently increased (Supplementary information, Fig. S2b). In another aforementioned CRISPR–Cas9 screen that was performed in monoclonal HUDEP-2 cells (GM-Clone 13) with homozygous mutations in the TGACCA motif, sgRNAs that targeted CACCC motifs in the *HBB* promoter were also enriched (Supplementary information, Fig. S2c and Table S1). We also obtained monoclonal HUDEP-2 cells in which both motifs (four copies of *HBG* and two copies of *HBB*) were edited, and we observed that γ-globin was almost exclusively expressed, while β-globin expression was nearly absent in these cells, achieving near complete reversal of the γ-to β-globin switch (Fig. 2c, d; Supplementary information, Table S1). Furthermore, mutating both motifs enhanced the *HBG*-LCR interaction and increased the transcriptional bursting frequency of *HBG*, whereas the *HBB*-LCR interaction was weakened and the transcriptional bursting frequency of *HBB* was decreased (Fig. 2e, f; Supplementary information, Fig. S3 and Table S1). We also observed that mutating both motifs increased chromatin accessibility, RNAPII binding and the active transcriptional epigenetic modification H3K4me3 and H3K9Ac on *HBG*, while decreased that on *HBB*, accompanied by changes in the binding of transcription factor complexes to the β-globin gene cluster (Fig. 2g; Supplementary information Fig. S2d, S3). These results indicate that further relieving the epigenetic repression of *HBG* after disrupting the KLF1-mediated *HBB*-LCR interaction reinforces the interaction between *HBG* and the LCR, thereby near completely reversing the γ-to β-globin switch.

**Fig 2.**
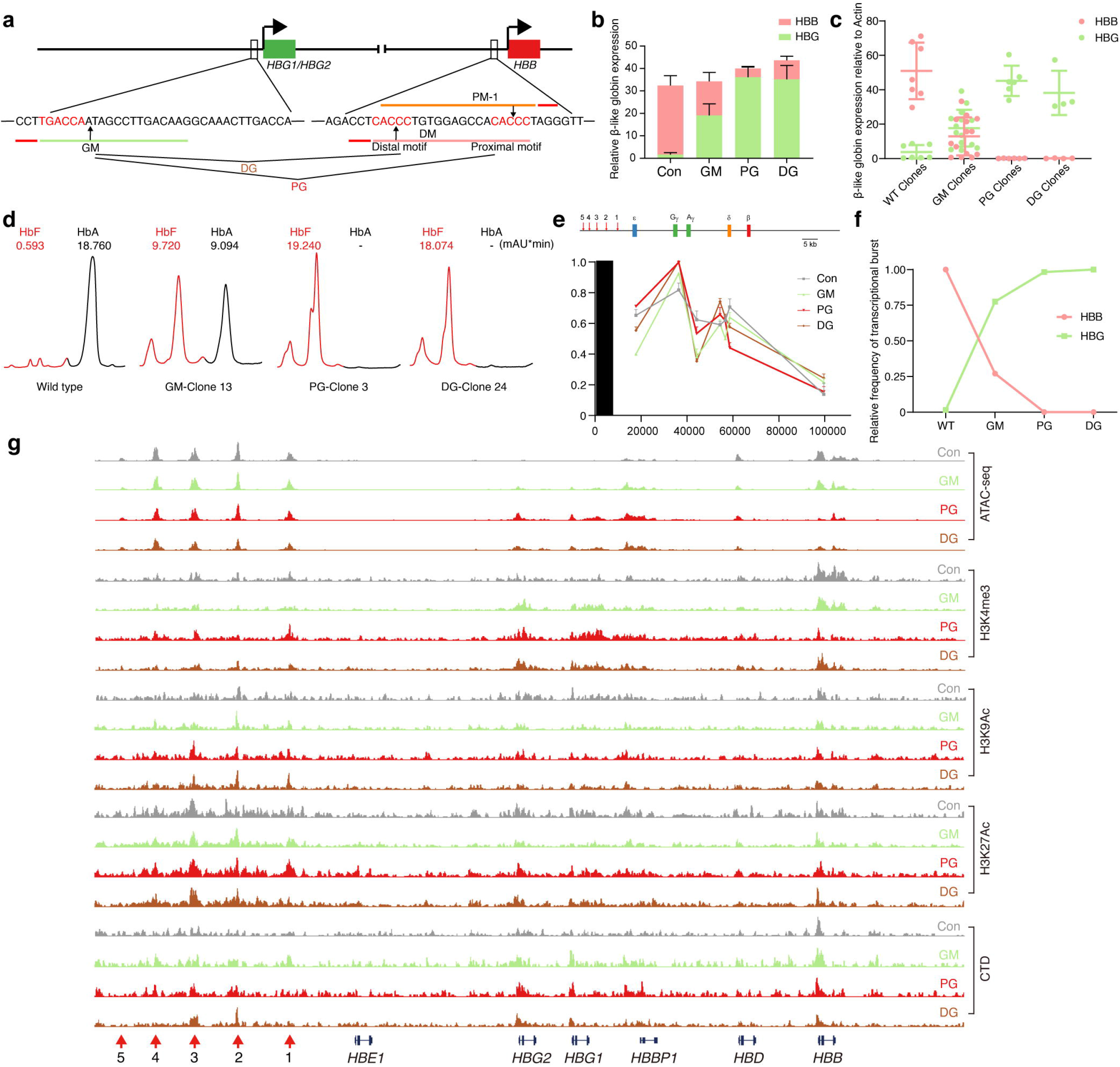
Mutations in the CACCC motif in *HBB* and the TGACCA motif in *HBG* near completely reverse the γ-to β-globin switch. **a** Cas9-expressing HUDEP-2 cells were transfected with sgRNAs targeting the TGACCA motif in *HBG*, followed by subsequent transfection with sgRNAs targeting the CACCC motif in *HBB*. **b** β-like globin gene expression relative to β-actin mRNA expression in HUDEP-2 cells from (**a**) as measured by RT–qPCR (mean ± s.d., n = 3). **c** β-like globin mRNA levels relative to β-actin mRNA levels in HUDEP-2 wild type (WT) clones (n = 7), GM clones (n = 14), PG clones (n = 6) and DG clones (n = 4) on day 5 of erythroid differentiation (mean ± s.d.). All GM, PG and DG clones carried four copies of *HBG*. **d** Fetal hemoglobin protein levels (normalized to total protein at 280 nm per 100 mAU*min) in four clonal populations from (**c**) as determined by HPLC. **e** Fetal Chromosome conformation capture analysis of HUDEP-2 cells from (**a**) (mean ± s.d., n = 3). **f** The relative frequency of the transcriptional bursts of *HBB* and *HBG* in HUDEP-2 clones from (**c**) were assessed by Chromium single cell sequencing. **g** ATAC-seq signals at the β-globin cluster were analyzed in HUDEP-2 cells from (**a**), along with CUT&Tag enrichment for CTD, H3K4me3, H3K9Ac, and H3K27Ac.

### Near completely reversing the γ-to β-globin switch in vivo

To confirm the effects of simultaneous mutation of the CACCC motif and TGACCA motif on the globin switch in vivo, the β-BAC mouse line (β D3), which is a transgenic mouse carrying a single copy of the ∼97 kb human β-globin locus that was generated in our laboratory^35^, was utilized to establish mouse lines with mutations of the TGACCA motif alone (GM-BAC) or mouse lines with simultaneous mutations of the CACCC motif and TGACCA motif (PG-BAC, DG-BAC) (Fig. 3a). Hemizygotes of these mouse lines were analyzed for globin expression at different stages of development from embryonic day 10.5 to the neonatal stage. Wild-type β-BAC mice exhibited a normal γ-to β-globin switch, while GM-BAC mice displayed a markedly delayed globin switch, with γ-globin expression still suppressed in the neonatal stage. In contrast, in both PG-BAC and DG-BAC mice, γ-globin was almost exclusively expressed and β-globin was nearly absent from embryonic day 12.5 to the neonatal stage, indicating that simultaneous mutation of the CACCC motif in *HBB* and the TGACCA motif in *HBG* near completely reversed the γ-to β-globin switch in vivo (Fig. 3b).

**Fig 3.**
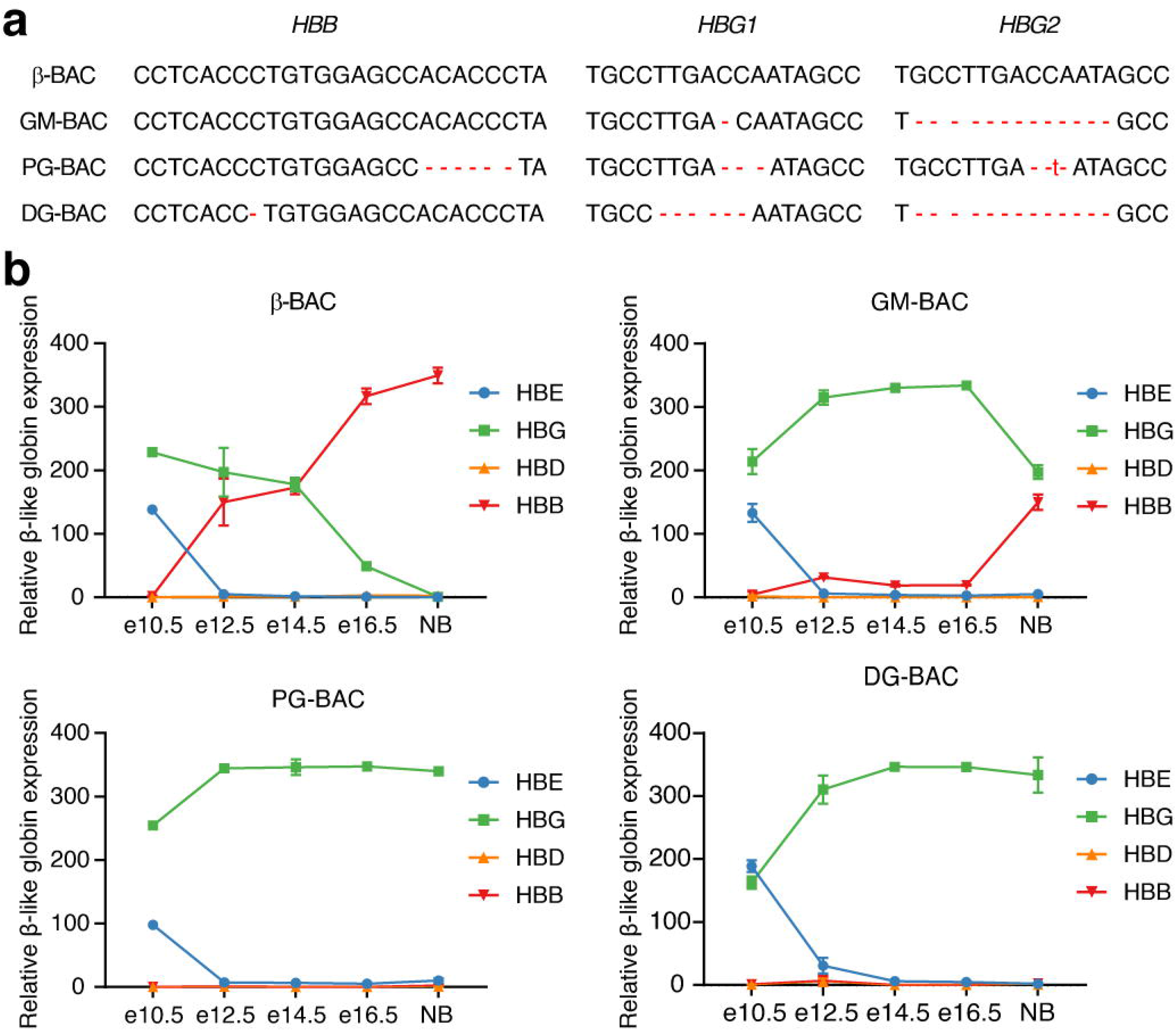
Near completely reversing the γ-to β-globin switch in β-BAC mice. **a** The β-BAC mouse line and three BAC mouse lines with mutations in the TGACCA motif alone (GM-BAC) or with mutations in both the CACCC motif and TGACCA motif (PG-BAC and DG-BAC) were established. The *HBB, HBG1* and *HBG2* genotypes corresponding to each line are shown. **b** Expression of human β-like globin genes relative to mouse β-actin mRNA expression was measured by RT–qPCR in embryonic day 10.5 (e10.5) yolk sacs, e12.5, e14.5 and e16.5 fetal livers, and newborn spleens of β-BAC, GM-BAC, PG-BAC and DG-BAC mice (mean ± s.d., n ≥ 3).

### Combined editing of the CACCC and TGACCA motifs increases HbF expression in human primary erythroblasts

Elevating the HbF level can attenuate hemoglobin polymerization and thus ameliorate the pathology of β-hemoglobinopathies^4,5^. To test whether simultaneous editing of the CACCC and TGACCA motifs can induce HbF expression in human primary erythroblasts, we first transfected normal peripheral blood-mobilized human donor CD34^+^ HSPCs with RNP complexes and then induced erythroid differentiation in vitro. Compared with editing the TGACCA motif alone, editing both the CACCC and TGACCA motifs increased γ-globin expression by 1.5-fold (PG) or 1.2-fold (DG) (fig. S4A). The HbF level increased from 79.3 mAU*min to 120.2 mAU*min and 91.2 mAU*min, respectively (Fig. 4a). The proportion of F cells also consistently increased (Supplementary information, Fig. S4b). Moreover, editing in CD34^+^ cells did not alter erythroid differentiation, as evaluated by flow cytometry (Fig. 4b). We then performed genome editing in CD34^+^ cells from patients with β^0^-thalassemia (codon 17 (A>T)/codon 41/42 (–TTCT)). Consistent with the results from HUDEP-2 cells and normal primary adult erythroblasts, simultaneous editing of these two motifs strongly induced γ-globin expression and increased the proportion of F cells (Fig. 4c-e; Supplementary information Fig. S5a). We also obtained erythroid clones that differentiated from single CD34^+^ cells in which both motifs were edited, and we observed a 1.5-fold increase in γ-globin expression compared with clones in which only the TGACCA motif was edited (Supplementary information, Fig. S5b and Table S2). In addition, motif editing in CD34^+^ cells from patients with β^0^-thalassemia did not affect erythroid differentiation (Supplementary information, Fig. S5c). Thus, combined editing of the CACCC and TGACCA motifs increased γ-globin expression in human primary erythroblasts.

**Fig 4.**
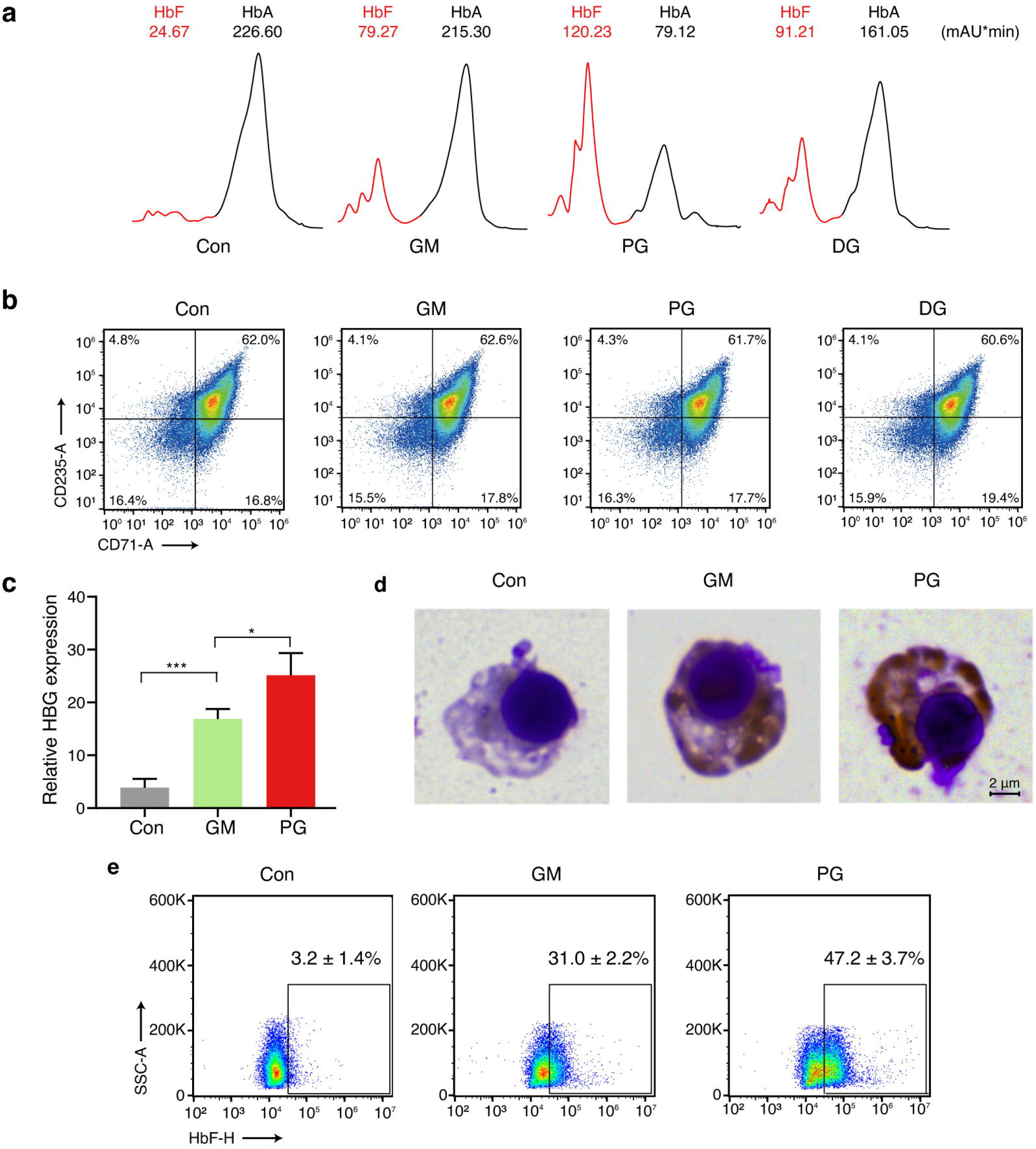
Combined disruption of the CACCC and TGACCA motifs increases HbF expression in primary erythroblasts. **a** Normal CD34^+^ HSPCs were transfected with RNPs targeting the distal TGACCA motif in *HBG*, followed by subsequent transfection of RNPs targeting the CACCC motifs in *HBB* and induction of erythroid differentiation. The average editing efficiency of *HBG* was 73.4%, and that of *HBB* in PG and DG was 76.4% and 11.6%, respectively. Fetal hemoglobin protein levels (normalized to total protein at 280 nm per 100 mAU*min) were determined by HPLC on day 12 of erythroid differentiation. **b** Flow cytometry analysis of the erythroid maturation markers CD71 and CD235 in the cells from (**a**). **c** CD34^+^ HSPCs from β^0^-thalassemia patients (codon 17 (A>T)/codon 41/42 (–TTCT)) were transfected with RNPs targeting the distal TGACCA motif in *HBG*, followed by subsequent transfection of RNPs targeting the proximal CACCC motifs in *HBB* and induction of erythroid differentiation. The average editing efficiency of *HBG* was 76.1%, and that of *HBB* in PG was 73.7%. The chart shows γ-globin gene expression relative to β-actin mRNA expression as measured by RT–qPCR on day 12 of erythroid differentiation (mean ± s.d., n = 2). Multiple comparisons were assessed with one-way ANOVA with Tukey’s MCT. \**P* < 0.05, \*\*\**P* < 0.001. **d** Hemoglobin accumulation as determined by benzidine– Giemsa staining in cells from (**c**) after 12 days of erythroid differentiation. **e** Representative flow cytometry plots showing HbF^+^ erythroblasts, derived as described in (**c**).

## Discussion

After completion of the γ-to β-globin switch in adult, the LCR interacts with *HBB* to activate its expression, while *HBG* is epigenetically silenced^1,9,24^ (Fig. 5a). We mutated the CACCC motif in *HBB* to disrupt the interaction between *HBB* and the LCR, resulting in the release of the LCR from *HBB* and its retargeting to other β-like globin genes, accompanied by a modest increase in other β-like globin (Fig. 5b). Subsequently, we simultaneously mutated the TGACCA motif in *HBG* and the CACCC motif in *HBB* to further relieve the epigenetic repression of *HBG*, thereby reinforcing the interaction between *HBG* and the LCR, leading to the expression of HbF at nearly 100% of the total hemoglobin level, thus near completely reversing the γ-to β-globin switch (Fig. 5c). This finding demonstrated that silenced genes can be fully activated through the redirection of enhancer-promoter contacts.

**Fig 5.**
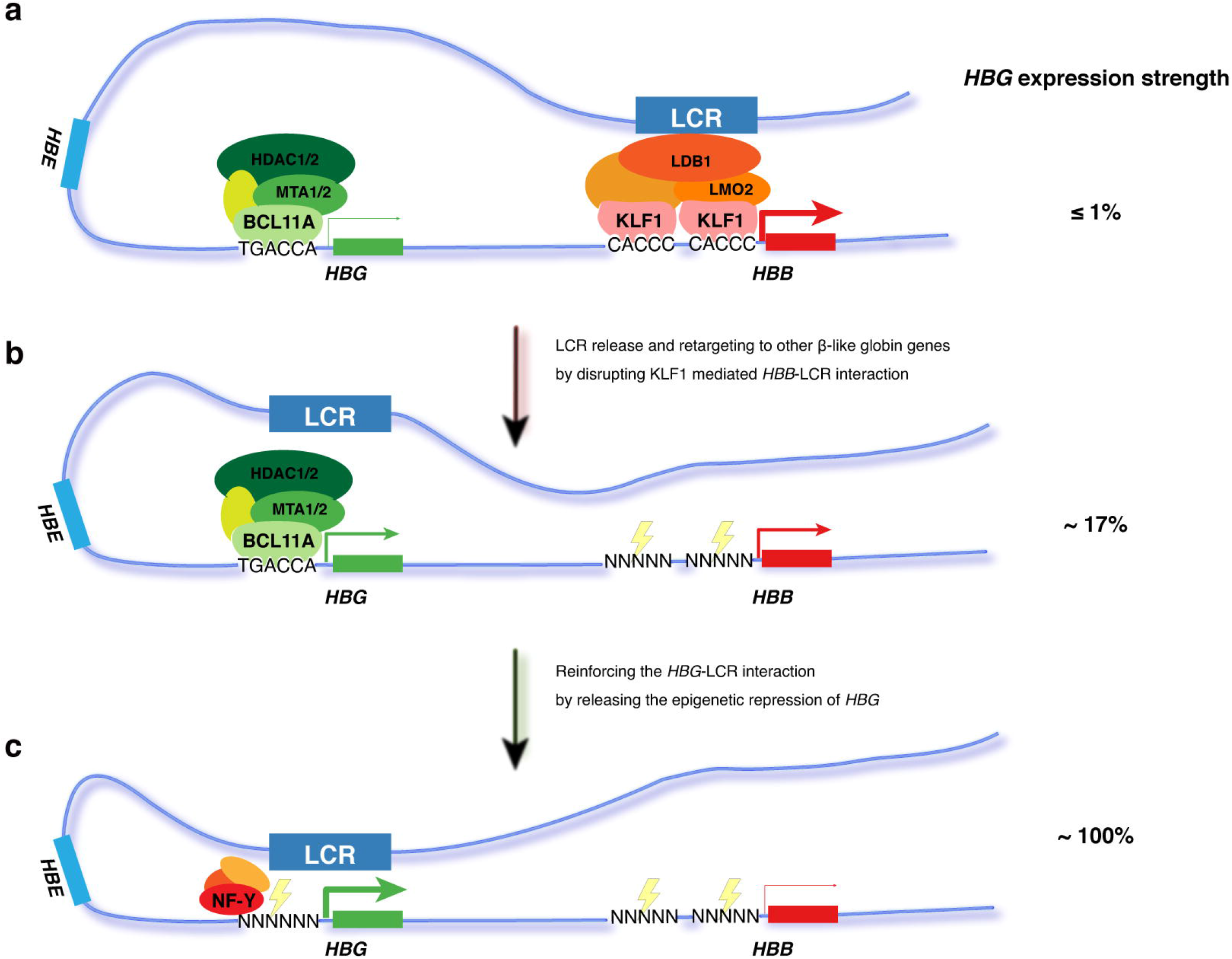
Diagram of near completely reversing the γ-to β-globin switch by LCR release, retargeting and reinforcing. **a** In adult erythrocytes, KLF1, along with the LMO2 complex and other factors, cooperatively mediates the spatial proximity between *HBB* and the LCR, activating β-globin expression. On the other hand, BCL11A, together with the NuRD complex and other factors synergistically suppresses *HBG* by epigenetic repression, resulting in γ-globin expression levels below 1%. **b** Mutating the CACCC motif disrupts the KLF1-mediated interaction between *HBB* and the LCR, leading to LCR release from *HBB* and retargeting to *HBG*, accompanied by increased γ-globin expression; however, *HBG* remains suppressed by epigenetic repression. **c** Simultaneously mutating the TGACCA motif and CACCC motif further releases the epigenetic repression of *HBG* controlled by BCL11A to reinforce the interaction between *HBG* and the LCR, leading to nearly 100% reactivation of *HBG* and near completely reversal of the γ-to β-globin switch.

The γ-to β-globin switch is a classic research model of eukaryotic gene expression regulation, and previous studies have identified numerous regulators involved in the activation of β-globin expression and the silencing of the γ-globin gene in adulthood^6,9,23,27^. However, these two aspects of the globin switch mechanism have only been studied separately, resulting in a lack of a comprehensive explanation for γ-to β-globin switching. Notably, a previous study on deletional HPFH focused on the *HBB* promoter, where the disruption of its promoter activity led to activation of *HBG*, providing the first experimental evidence that activation of *HBB* in adult also contributes to the silencing of *HBG*^*15*^. Here, we employed CRISPR-Cas9 screening and identified the major *HBB* activator KLF1’s CACCC motif on *HBB* promoter participates the repression of *HBG* in adults. More importantly, for the first time, we achieved near complete reversal of the γ-to β-globin switch by simultaneously mutating the CACCC motif in *HBB* and the TGACCA motif in *HBG*, suggesting that the γ-to β-globin switch is comprehensively regulated by the epigenetic silencing of *HBG* and the competitive repression between *HBB* and *HBG*.

Enhancer-promoter engagement ensures spatiotemporal gene expression^20,36^. A previous study has shown that this specificity between enhancer and its target promoter is determined by the CTCF binding at promoters^21^. However, simultaneously mutating the binding sites of two transcription factors is sufficient to near completely redirect the LCR from interacting with *HBB* to interacting with *HBG*. This finding indicates that the specificity of enhancer-promoter contact within chromosomal domains is achieved through the binding of transcription factor clusters to enhancers and promoters, providing experimental evidence for enhancer-promoter specification in the ‘selecting-facilitating-specifying’ model^20^. Additionally, these results indicate that different transcription factor clusters exhibit compatibility with the same enhancer, which underlies the contribution of spatial conformation competition to gene silencing when two or more promoters are controlled by the same enhancer. Thus, disrupting one gene promoter can increase the expression of another gene that is regulated by the same enhancer^15,21^. Furthermore, previous research has indicated that transcriptional bursting frequency is primarily determined by enhancers^31,32^. Our data, however, show that the attributes of promoters also contribute to the determination of bursting frequency.

β-Hemoglobinopathies are the most common monogenic genetic diseases worldwide^2,3^. Manipulating hematopoietic stem cells via genome editing technology has promoted innovative strategies for treating β-hemoglobinopathies^37^. Currently, the most appropriate gene therapy strategy for activating γ-globin expression involves targeting the core sequence in the erythroid-specific enhancer of BCL11A or the TGACCA motif of BCL11A and other HPFH-related point mutations in *HBG*^38-44^. Manipulating these sites can theoretically increase the HbF level up to approximately 60% of the total hemoglobin level, ameliorating the clinical symptoms of β-hemoglobinopathies^38,45^. However, in populations with HPFH, variations in disease severity indicate that β-hemoglobinopathy patients with this level of HbF still exhibit certain disease symptoms, and as HbF levels further increase, the clinical manifestations of patients become milder, necessitating further increases in the HbF levels^16,17^. Combined editing of the CACCC motif in *HBB* and the TGACCA motif in *HBG* can elevate HbF level to nearly 100% of the total hemoglobin level, suggesting that this approach may be a more optimal gene therapy strategy for β-hemoglobinopathies. And a single delivery of base editing system carrying two different sgRNAs is more suitable for the clinical application of this gene therapy strategy, which requires simultaneous editing of CACCC motif in *HBB* and the TGACCA motif in *HBG* within one chromosome. Furthermore, the simultaneous disruption of epigenetic silencing and spatial conformation competition also shows potential for application in other scenarios, such as reactivating suitable genes for tissue regeneration or tumor suppressor genes for cancer therapy^46-48^.

## Materials and Methods

### HUDEP-2 cell culture and induction of erythroid differentiation

HUDEP-2 cells were obtained from R. Kurita and Y. Nakamura (RIKEN BioResource Center) and expanded in StemSpan serum-free expansion medium (SFEM, Stem Cell Technologies) supplemented with 1 μM dexamethasone, 1 μg/ml doxycycline, 50 ng/ml SCF (PeproTech), and 3 U/ml EPO (PeproTech). Cells were differentiated for 5 d in Iscove’s modified Dulbecco’s medium (IMDM) supplemented with 330 μg/ml holo-transferrin (Sigma), 3 IU/ml heparin, 10 μg/ml insulin, 5% AB human serum (Gemini), 3 IU/ml EPO, and 1 μg/ml doxycycline. The cell density was maintained between 0.7 × 10^6^/ml and 1.4 × 10^6^/ml for the first three days and then between 1 × 10^6^/ml and 2 × 10^6^/ml for the following days.

### Human primary CD34^+^ HSPC culture and in vitro erythroid differentiation

Normal human CD34^+^ HSPCs from mobilized peripheral blood were obtained from Hycells, and β-thalassemia CD34^+^ HSPCs from bone marrow were obtained from three β^0^-thalassemia donors (codon 17 (A>T)/codon 41/42 (–TTCT)) at the First Affiliated Hospital of Guangxi Medical University. Human CD34^+^ cells were expanded in SFEM supplemented with 100 ng/mL SCF, 100 ng/mL FLT3-L, and 100 ng/mL TPO. For erythroid differentiation, CD34^+^ HSPCs were first cultured in IMDM supplemented with 5% human AB serum, 200 μg/mL holo-human transferrin, 10 μg/mL insulin, 3 IU/mL heparin, 10 ng/mL SCF, 1 ng/mL IL-3, and 3 IU/mL EPO for 7 days, followed by an additional 5 days of differentiation in medium without IL-3. Erythroid differentiation was monitored by flow cytometry using PE-conjugated anti-CD71 (BioLegend, 334108) and FITC-conjugated anti-CD235 antibodies (BioLegend, 349104).

### CRISPR-Cas9 screen for regulatory motifs that modulate γ-globin expression

A total of 3,784 sgRNAs were designed targeting the downstream region of *HBG* (hg38/GRCh38, chr11: 5204054-5248601) ^49^, followed by the generation of a pooled lentiviral vector library using the lentiGuide-Puro plasmid (Addgene, plasmid 52963) ^50^ that contains a puromycin-resistance cassette for selection. Approximately 1.2×10^7^ HUDEP-2 cells stably expressing Cas9 were transduced with the lentiviral library at a multiplicity of infection of 0.3. After 24 hours, the infected cells were selected with 1 μg/ml puromycin for 3 days and subsequently cultured for an additional 6 days. Then, 1×10^7^ HUDEP-2 cells were collected as the control, and positive cells with the 2% highest anti-HbF (Invitrogen, MHFH05) staining intensities were purified from the remaining cells by FACS. Genomic DNA was extracted and the sgRNA library was prepared by PCR amplification. All PCR primers include the following: CRSIPR-Cas9 screening first step PCR primer (F: TCTTTCCCTACACGACGCTCTTCCGATCTAATGGACTATCATATGCTTACCGTAACTTGAAAGTATTTCG; R: AGTTCAGACGTGTGCTCTTCCGATCCTTTAGTTTGTATGTCTGTTGCTATTATGTCTACTATTCTTTCC), NGS second step PCR primer (F: AATGATACGGCGACCACCGAGATCTACACTCTTTCCCTACACGAC; R: CAAGCAGAAGACGGCATACGAGATNNNNNNGTGACTGGAGTTCAGACGTGTGCTCT TC). MiSeq 300-bp paired-end sequencing (Illumina) was performed. Candidate motifs with HbF regulatory function were identified by analyzing the sequencing data using CRISPR-SURF ^51^.

### Genome editing

Chemically modified single guide RNAs (PM-1: ACCCTGTGGAGCCACACCCT; PM-2: GATTGGCCAACCCTAGGGTG; DM: GGGTGTGGCTCCACAGGGTG; GM: CTTGTCAAGGCTATTGGTCA) were synthesized by Integrated DNA Technologies. For combined editing of the CACCC motif and TGACCA motif, cells were first electroporated with RNP complexes containing GM and were then expanded for 36 hours. The subset of edited cells was transduced with RNP complexes containing PM or DM in the second electroporation. Due to the low editing efficiency of DM, additional electroporation of RNP complexes containing DM was performed in the DM or DG groups of HUDEP-2 cells. For transient editing, the Neon Transfection System was used to electroporate HUDEP-2 cells (1200 V, 40 ms, 1 pulse), and the Lonza 4D-Nucleofector system was used to electroporate CD34^+^ HSPCs (E0-100, solution P3). Editing efficiency was analyzed by CRISPResso2 ^52^ after NGS sequencing for PCR amplificon of the genome target. All PCR primers include the following: NGS-HBB (F: TCTTTCCCTACACGACGCTCTTCCGATCTTAGCCAGTGCCAGAAGAGC; R: AGTTCAGACGTGTGCTCTTCCGATCCACCACCAACTTCATCCAC), NGS-HBG (F: TCTTTCCCTACACGACGCTCTTCCGATCTGGAATGACTGAATCGGAACA; R: AGTTCAGACGTGTGCTCTTCCGATCTGGAACTGCTGAAGGGTG) and NGS second step PCR primer. After editing, monoclonal cells were obtained and genotyped as indicated in table S1 for HUDEP-2 cells and in table S2 for CD34^+^ HSPCs.

### Experimental animals

The β-globin locus transgenic (β-BAC) mouse strain (β D3) was established by microinjection of a BAC containing the ∼97 kb human β-globin locus. To obtain mouse strains with mutations in both the CACCC motif and the TGACCA motif, RNPs containing PM-1 (or DM) and GM were microinjected into zygotes of β-BAC mice. GM-BAC, PG-BAC and DG-BAC mice were identified and maintained in the hemizygous state. Ter119-positive erythrocytes were isolated from the embryonic 10.5 (e10.5) yolk sacs, e12.5, e14.5 and e16.5 fetal livers and newborn spleens of these transgenic mice. Total RNA was extracted for quantification of globin mRNA. All animal procedures were approved by the Animal Care and Use Committee at the Institute of Basic Medical Sciences, Chinese Academy of Medical Sciences, and Peking Union Medical College.

### Real-time qPCR

Total RNA was extracted from 0.5-2 million cells according to the protocol of TRIzol (Invitrogen). Reverse transcription was performed using the PrimeScript RT Reagent Kit (Takara). mRNA was quantified by SYBR Green qPCR. All PCR primers include the following: HBE (F: GCAAGAAGGTGCTGACTTCC; R: ACCATCACGTTACCCAGGAG), HBG (F: TGGATGATCTCAAGGGCAC; R: TCAGTGGTATCTGGAGGACA), HBD (F: GAGGAGAAGACTGCTGTCAATG; R: AGGGTAGACCACCAGTAATCTG), HBB (F: CTGAGGAGAAGTCTGCCGTTA; R: AGCATCAGGAGTGGACAGAT), Mouse β-actin (F: ACCCTAAGGCCAACCGTGA; R: GTCTCCGGAGTCCATCACAA), Human β-actin (F: CACCCAGCACAATGAAGATC; R: GTCATAGTCCGCCTAGAAGC).

### Hemoglobin HPLC

Approximately 2×10^6^ cells were harvested and washed with PBS. Cells were lysed in 20 μl of 0.01% SDS and incubated for 10 min on ice. The lysate was then filtered through Ultrafree-MC filters (PVDF, 0.45 µm) and incubated in 80 μl buffer A (20 mM Bis-Tris, pH 6.8) for 30 min on ice. Hemoglobins were eluted from the PolyCAT A column (PolyLC, 3.54CT0315) during a 2% to 25% gradient of buffer B (20 mM Bis-Tris, 200 mM NaCl, pH 6.9) in buffer A for 10 min followed by a 25-100% gradient of buffer B in buffer A for 10 to 22 min at a flow rate of 1.2 ml/min. Hemoglobin proteins were detected by absorbance measurements at 415 nm and normalized to total protein at 280 nm per 100 mAU*min.

### HbF flow cytometry analysis

Cells were fixed with 0.05% glutaraldehyde, permeabilized with 0.1% Triton X-100, and stained with anti-human HbF-APC for flow cytometry according to the manufacturer’s recommendations. Analytical flow cytometry was performed using a Beckman Coulter CytoFLEX Cytometer.

### Chromosome Conformation Capture Analysis

Approximately 10^7^ cells were subjected to crosslinked with 1% formaldehyde for 10 min, and the reaction was then quenched with glycine. Cross-linked cells were lysed for 30 min with cold lysis buffer (10 mM Tris-HCl at pH 8.0, 10 mM NaCl, 0.2% NP-40), resuspended in 1× NEB CutSmart buffer with 0.1% SDS, and then shaken for 5 min at 65°C. Triton X-100 was added to 2%, and the nuclei were shaken for 1 hour at 37°C. Digestion was carried out using EcoRI-HF at a final concentration of 2 U/μl, followed by ligation using DNA ligase at a final concentration of 0.25 U/μl. After decrosslinking, the DNA was purified by phenol extraction and isopropanol precipitation and quantified by qPCR. All PCR primers include the following: 3C-HS2-4: ACATAGTTGTCAGCACAATGCCTA, 3C-HS1: TTTTAGCACTTACAGTCTGCCAAA, 3C-HBG2: TATTGTCTCCTTTCATCTCAACAGC, 3C-HBG+3k: GAGCGGGTGAGAGAAAAGTG, 3C-DELTA: GGGTGTGTATTTGTCTGCCA, 3C-HBD: CATGTATCTGCCTACCTCTTCTCC, 3C-HBB: ATAGTAATTGTGTGTGCTCGGC, 3C-3Distal: CACCAACTCCCAGCTTTACTAACT. Samples were normalized to an independent ERCC3 locus.

### Single-cell Multiome ATAC + Gene Expression

Single-cell Multiome ATAC + Gene Expression was performed with Chromium Single Cell Multiome ATAC + Gene Expression (10× Genomics) in HUDEP-2 clones (Wild type, PM-Clone 22, DM-Clone 13, GM-Clone 13, PG-Clone 3, DG-Clone 24) following the manufacturer’s protocol. Raw sequencing data was processed with the Cell Ranger ARC pipeline (version 2.0.0) and mapped to the human reference genome hg38. The intranuclear *HBB*/*HBG* RNA levels were obtained after normalization and linear and non-linear dimension reduction and clustering by Seurat ^53^. Gene activity score of *HBB*/*HBG* was generated by analyzing the ATAC-seq signals on the 2 kb upstream region of *HBB/HBG* using Signac ^54^.

### ATAC-seq

Fifty thousand viable HUDEP-2 cells were lysed in Lysis Buffer (10 mM Tris-HCl at pH 7.4, 10 mM NaCl, 3 mM MgCl_2_, 0.1% Tween 20, and 0.01% Digitonin). Nuclear pellet was then subjected to transposition reaction using Hyperactive ATAC-Seq Library Prep Kit for Illumina (Vazyme). Cleaned up libraries were sequenced by MiSeq 300-bp paired-end sequencing (Illumina). ATAC-seq data were mapped to the human reference genome hg38 using bowtie 2 ^55^ and converted to bigwig files for visualization in UCSC browser.

### CUT&Tag

Fifty thousand viable HUDEP-2 cells were used to generate libraries using a NovoNGS CUT&Tag 4.0 High-Sensitivity Kit (Novoprotein) according to the manufacturer’s instructions. anti-RNA polymerase II CTD repeat YSPTSPS antibody-ChIP Grade(abcam, ab26721), anti-Histone H3(tri methyl K4) ChIP Grade(abcam, ab8580), anti-H3K9Ac ChIP Grade(abcam, ab4441), anti-Histone H3 (acetyl K27) ChIP Grade(abcam, ab4729), anti-GATA1(abcam, ab11852), anti-KLF1(abcam, ab2483), anti-LDB1(CST, 64994), anti-CHD4(CST, 12011), anti-MBD2(abcam, ab188474), anti-NFYA(abcam, ab139402) and anti-CTCF(CST, 3418) were used for CUT&Tag. Cleaned up libraries were sequenced by MiSeq 300-bp paired-end sequencing (Illumina). ATAC-seq data were mapped to the human reference genome hg38 using bowtie 2 ^55^ and converted to bigwig files for visualization in UCSC browser.

### Benzidine-Giemsa staining

Cells deposited on glass slides were fixed in methanol for 4 min and stained with 3,3′-dimethoxy-benzidin and Giemsa stain following the instructions provided by Sigma Aldrich. The stained cells were examined and imaged by microscopy.

## Data and materials availability

The authors declare that all the data supporting the findings of this study are available within the article and supplementary materials. All materials in this article are available upon reasonable request from the corresponding authors.

## Acknowledgements

We thank Peng Xu at Soochow University for insightful discussions and technical support since the inception of this work in 2017. We thank Zemin Zhang at Peking University for helpful advice on the manuscript. We thank members of the Liu laboratory for helpful discussions and technical assistance. We thank State Key Laboratory of Common Mechanism Research of Major Diseases Platform for consultation and instrument availability that supported this work.

## Author Contributions

N.W. conceived and designed the experiments; N.W. performed the majority of experiments, assisted by K.Y., X.Z., S.C., X.P., and D.H.; K.Y. performed bioinformatics data analysis; X.Z. generated the transgene and edited mice; X.X, Y.J., G.Y., R.L., P.C., W.D., Y.H., and X.L. gave some helpful advice; N.W. and D.L. wrote the manuscript, with input from all authors; D.L. and Z.Z. supervised the study.

## Supplementary Materials

Figs. S1 to S5

Tables S1 to S2

