## Supplementary information, Fig. S1 for "Near completely reversing the γ- to β-globin switch by enhancer release, retargeting and reinforcing"

a

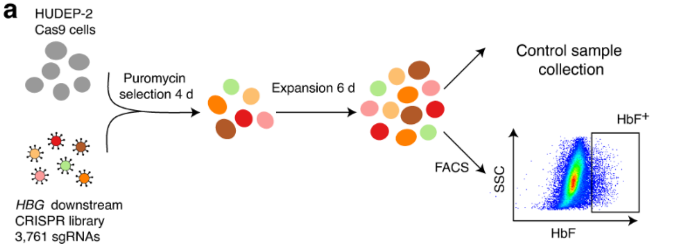


b

| Rank | gRNA sequence | Position | Targets |
|---|---|---|---|
| 1 | GCACTGTGACAAGCTGCACG | 5234014<br>5226600 | homologous sequence of *HBB* and *HBD* |
| 2 | CATGGTGCATCTGACTCCTG | 5234418<br>5227006 | homologous sequence of *HBB* and *HBD* |
| 3 | CTCAGGAGTCAGATGCACCA | 5234435<br>5227023 | homologous sequence of *HBB* and *HBD* |
| 4 | GGCTCACCTGGACAACCTCA | 5234062<br>5226648 | homologous sequence of *HBB* and *HBD* |
| 5 | ACCTTGATACCAACCTGCCC | 5234350<br>5226938 | homologous sequence of *HBB* and *HBD* |
| 6 | GCTCACCTGGACAACCTCAA | 5234061<br>5226647 | homologous sequence of *HBB* and *HBD* |
| 7 | TTGGTGGTGAGGCCCTGGGC | 5234347<br>5226935 | homologous sequence of *HBB* and *HBD* |
| 8 | GATTGGCCAACCCTAGGGTG | 5227159 | Proximal CACCC on *HBB* |
| 9 | TGGTGAGGCCCTGGGCAGGT | 5226931<br>5234343 | homologous sequence of *HBB* and *HBD* |
| 10 | ATGAAATTGGCCGGGCGCGG | 5244459 | BGLT3 |
| 11 | ACCCTGTGGAGCCACACCCT | 5227159 | Proximal CACCC on *HBB* |
| 12 | GCCCATAACAGCATCAGGAG | 5234157<br>5226743 | homologous sequence of *HBB* and *HBD* |
| 13 | TAGTGATGGCCTGGCTCACC | 5234074<br>5226660 | homologous sequence of *HBB* and *HBD* |
| 14 | GGTAGACCACCAGCAGCCTA | 5226805 | *HBB* intron1 |
| 15 | GTGTGGCAAAGGTGCCCTTG | 5226649 | *HBB* exon2 |
| 16 | TTAGAAAGTAGGGGTAGGAG | 5227982 | -900 *HBB* |
| 17 | AAGAAAGACCTCTATAGGAC | 5248015 | -2468 BGLT3 |
| 18 | AGAAAGTGCATATTCTGAAA | 5246777 | -1225 BGLT3 |
| 19 | GTGAGCCAGGCCATCACTAA | 5226680<br>5234094 | homologous sequence of *HBB* and *HBD* |
| 20 | ATCTTCCTCCCACAGCTCCT | 5225725 | *HBB* exon3 |

c

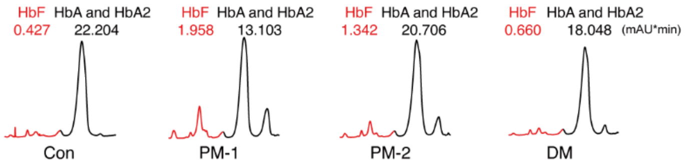


d

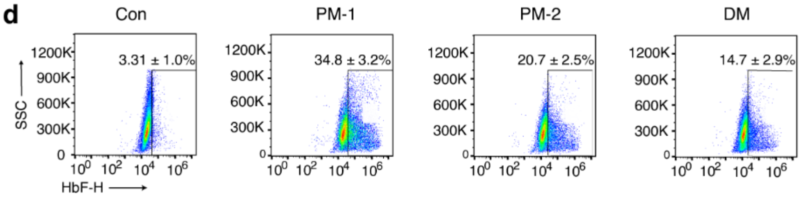


e

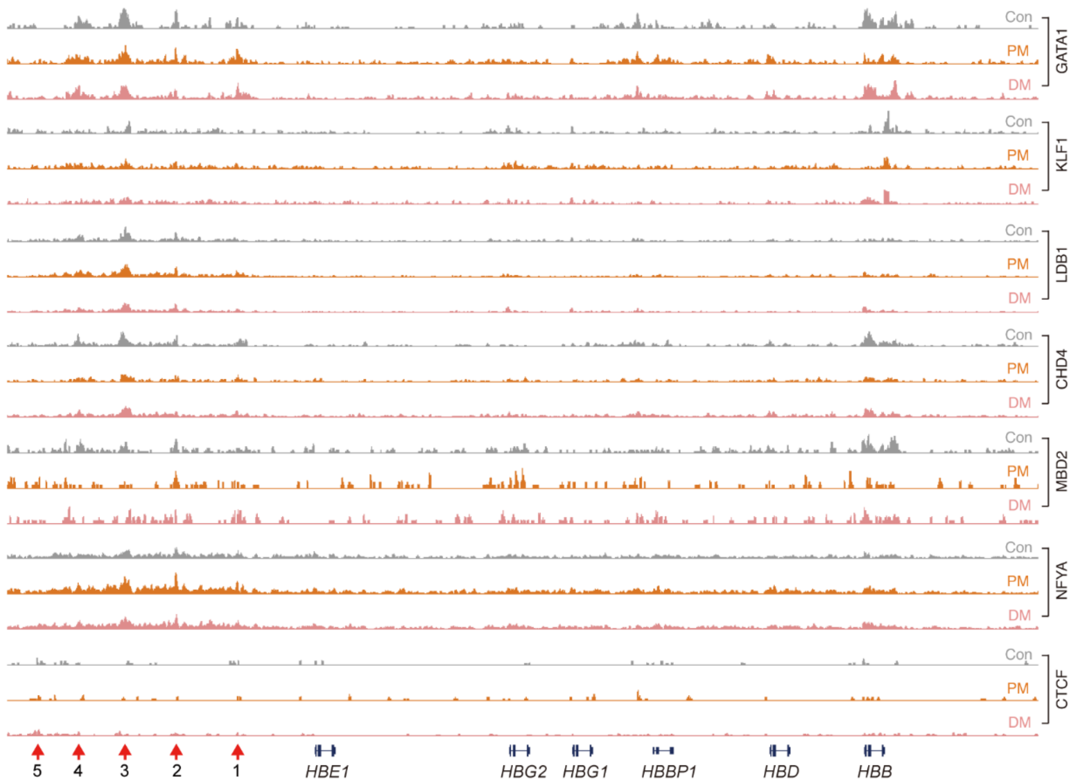


**Fig. S1 Induced HbF expression by disrupting the CACCC motif in *HBB* promoter. a** Workflow of CRISPR–Cas9 screening to identify regulatory elements of HbF. **b** The top 20 enriched sgRNAs in CRISPR–Cas9 screen. **c** Cas9-expressing HUDEP-2 cells were transduced with sgRNAs targeting the CACCC motif in *HBB*. Fetal hemoglobin protein levels (normalized to total protein at 280 nm per 100 mAU*min) were determined by high-performance liquid chromatography (HPLC) on day 5 of erythroid differentiation (n = 3). **d** Representative flow cytometry plots showing HbF$^{+}$ erythroblasts in control and CACCC motif edited HUDEP-2 cells (mean ± s.d., n = 3). **e** CUT&Tag signals at the β-globin cluster of GATA1, KLF1, LDB1, CHD4, MBD2, NFYA, and CTCF were analyzed in control and CACCC motif edited HUDEP-2 cells.
