## Supplementary information, Fig. S2 for "Near completely reversing the γ- to β-globin switch by enhancer release, retargeting and reinforcing"

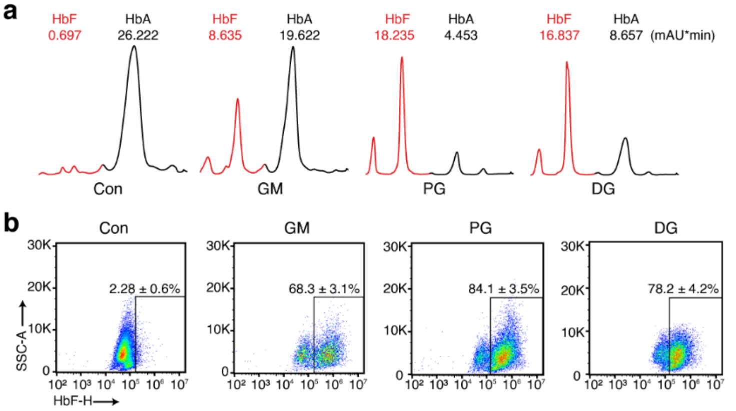


**c**

| Rank | gRNA sequence | Position | Targets |
|---|---|---|---|
| 1 | TTGGTGGTGAGGCCCTGGGC | 5234347<br>5226935 | homologous sequence of *HBB* and *HBD* |
| 2 | CATGGTGCATCTGACTCCTG | 5234418<br>5227006 | homologous sequence of *HBB* and *HBD* |
| 3 | ACCTTGATACCAACCTGCCC | 5234350<br>5226938 | homologous sequence of *HBB* and *HBD* |
| 4 | TGCTGTTATGGGCAACCCTA | 5234150<br>5226736 | homologous sequence of *HBB* and *HBD* |
| 5 | CCTTGATACCAACCTGCCCA | 5234351<br>5226939 | homologous sequence of *HBB* and *HBD* |
| 6 | ACCCTGTGGAGCCACACCCT | 5227159 | Proximal CACCC on *HBB* |
| 7 | ACGTGGATCCTGAGAACTTC | 5234016<br>5226602 | homologous sequence of *HBB* and *HBD* |
| 8 | AGTCTGCCGTTACTGCCCTG | 5226999 | *HBB* exon1 |
| 9 | GAGGTTCTTTGAGTCCTTTG | 5234189<br>5226775 | homologous sequence of *HBB* and *HBD* |
| 10 | GCCTATCAGAAAGTGGTGGC | 5225657 | *HBB* exon3 |
| 11 | CTCAGGAGTCAGATGCACCA | 5234435<br>5227023 | homologous sequence of *HBB* and *HBD* |
| 12 | CAGCTCACTCAGTGTGGCAA | 5226638 | *HBB* exon2 |
| 13 | GGCTGGTGTGGCTAATGCCC | 5233006<br>5225640 | homologous sequence of *HBB* and *HBD* |
| 14 | GCTCATGGCAAGAAAGTGCT | 5226708 | *HBB* exon2 |
| 15 | AGCACGTTGCCCAGGAGCTG | 5225732 | *HBB* exon3 |
| 16 | TGGTGAGGCCCTGGGCAGGT | 5226931<br>5234343 | homologous sequence of *HBB* and *HBD* |
| 17 | GAAGTTGGTGGTGAGGCCCT | 5226958 | *HBB* exon1 |
| 18 | CACAGACCAGCACGTTGCCC | 5225724 | *HBB* exon3 |
| 19 | TGATAGGCAGCCTGCACTGG | 5225669 | *HBB* exon3 |
| 20 | GATTGGCCAACCCTAGGGTG | 5227159 | Proximal CACCC on *HBB* |

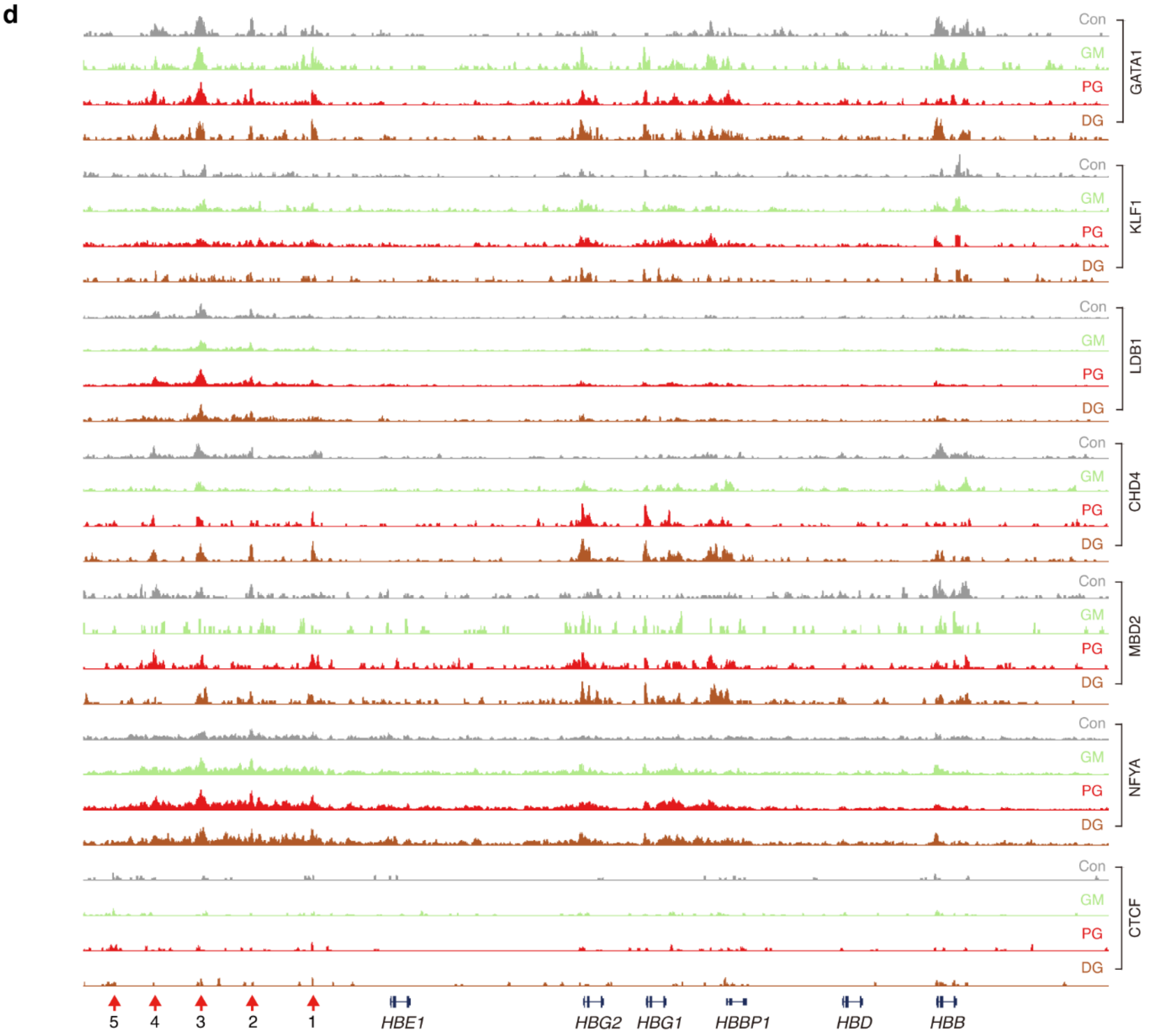


**Fig. S2 Mutations in the CACCC motif and TGACCA motif near completely reverse the γ- to β-globin switch. a** Cas9-expressing HUDEP-2 cells were transfected with sgRNAs targeting the TGACCA motif in *HBG*, followed by subsequent transfection with sgRNAs targeting the CACCC motifs in *HBB*. Fetal hemoglobin protein levels (normalized to total protein at 280 nm per 100 mAU*min) were determined by high-performance liquid chromatography (HPLC) (n = 3). **b** Representative flow cytometry plots showing $HbF^+$ erythroblasts among HUDEP-2 cells from (**a**) (mean ± s.d., n = 3). **c** The top 20 enriched sgRNAs in the CRISPR–Cas9 screen performed with GM-clone 13. **d** CUT&Tag signals at the β-globin cluster of GATA1, KLF1, LDB1, CHD4, MBD2, NFYA, and CTCF were analyzed in HUDEP-2 cells from (**a**).
