## Supplementary information, Fig. S3 for "Near completely reversing the γ- to β-globin switch by enhancer release, retargeting and reinforcing"

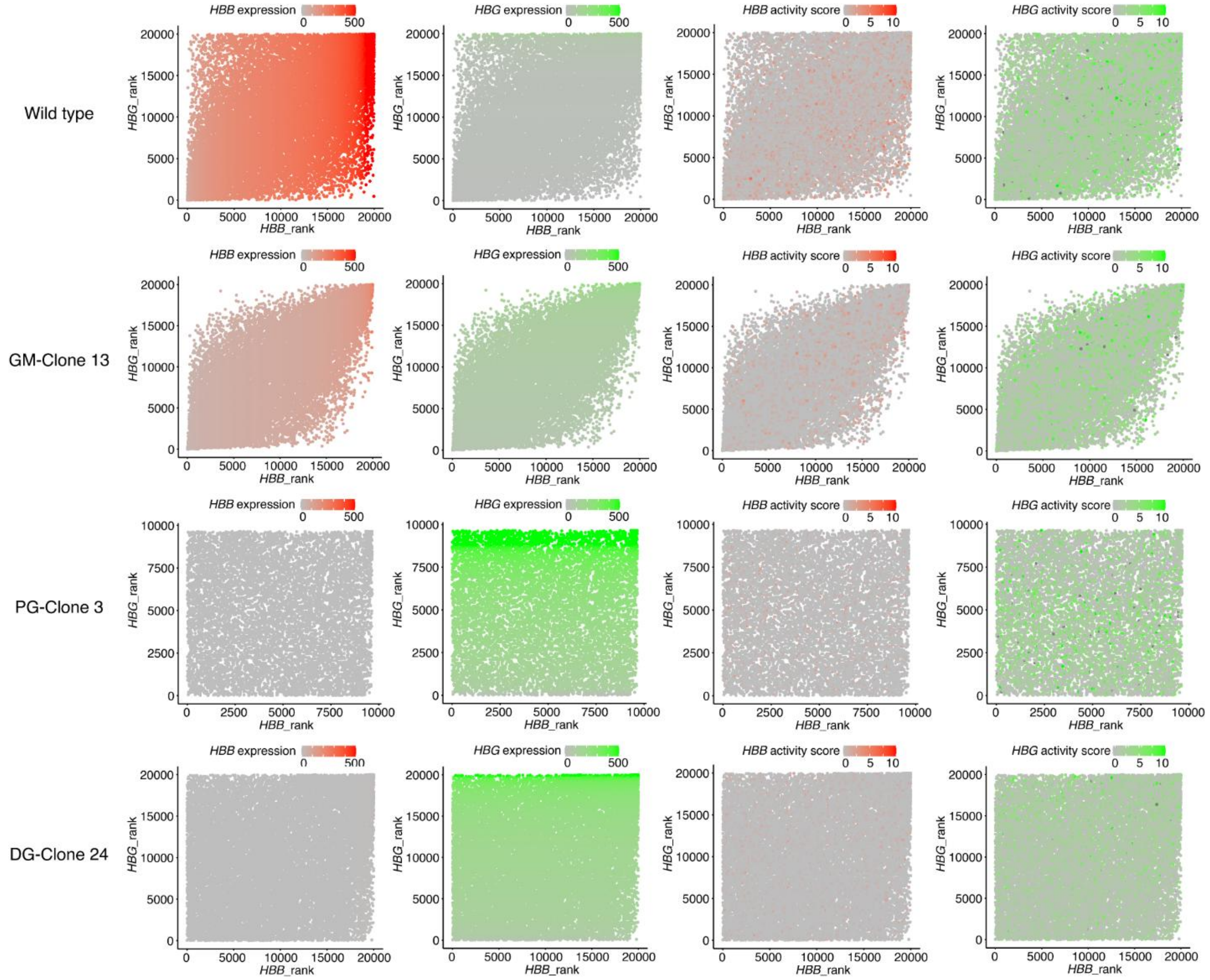


**Fig. S3 Single Cell Multiome ATAC + Gene Expression analysis of HUDEP-2 clones.**
The transcription and gene activity scores of *HBB* and *HBG* in HUDEP2 clones were analyzed by Chromium Single Cell Multiome ATAC + Gene Expression. The intranuclear *HBB*/*HBG* RNA levels represent gene expression, and the ATAC-seq signals at *HBB*/*HBG* indicate the gene activity score.
