## Supplementary information, Fig. S4 for "Near completely reversing the γ- to β-globin switch by enhancer release, retargeting and reinforcing"

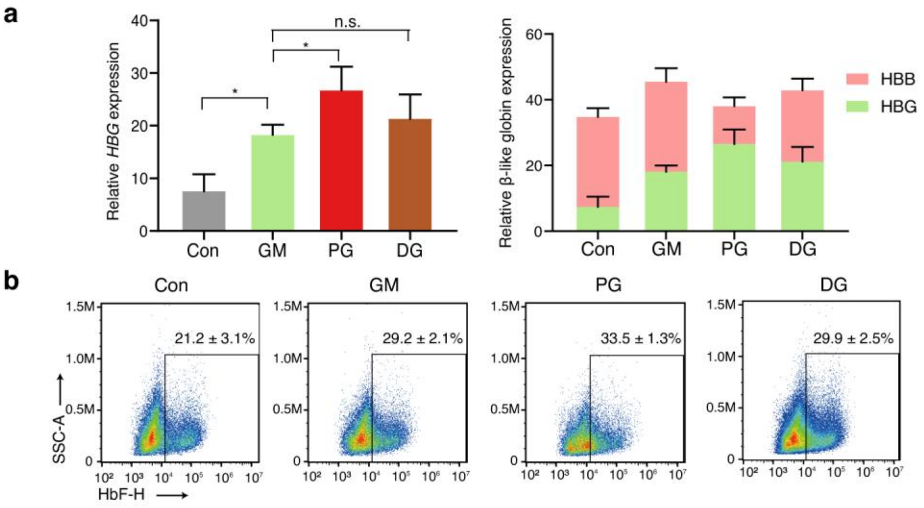


**Fig. S4 Combined disruption of the CACCC and TGACCA motifs increases γ-globin expression in human primary erythroblasts. a** Normal CD34$^+$ HSPCs were transfected with RNPs targeting the distal TGACCA motif in *HBG*, followed by subsequent transfection of RNPs targeting the CACCC motifs in *HBB* and induction of erythroid differentiation. The chart shows β-like globin gene expression relative to the β-actin mRNA expression as measured by RT–qPCR on day 12 of erythroid differentiation (mean ± s.d., n = 3). Multiple comparisons were assessed with one-way ANOVA with Tukey's MCT. n.s. not significant, $*P < 0.05$. **b** Representative flow cytometry plots showing HbF$^+$ erythroblasts derived as described in (**a**).
