## Supplementary information, Fig. S5 for "Near completely reversing the γ- to β-globin switch by enhancer release, retargeting and reinforcing"

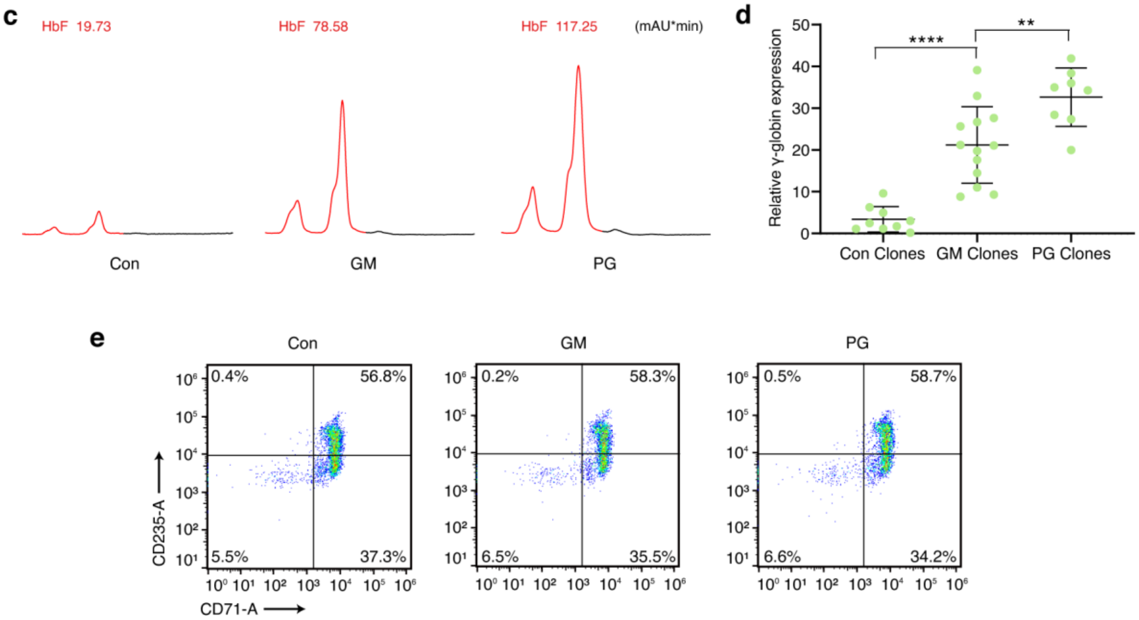


**Fig. S5 Combined disruption of the CACCC and TGACCA motifs stimulates HbF expression in HSPCs from $\beta^0$-thalassemia patients. a** $CD34^+$ HSPCs from $\beta^0$-thalassemia patients (codon 17 (A>T)/codon 41/42 (–TTCT)) were transfected with RNPs targeting the distal TGACCA motif in *HBG*, followed by subsequent transfection of RNPs targeting the proximal CACCC motifs in *HBB* and induction of erythroid differentiation. Fetal hemoglobin protein levels (normalized to total protein at 280 nm per 100 mAU*min) were determined by HPLC. **b** γ-globin mRNA levels relative to β-actin mRNA levels in $CD34^+$ WT clones (n = 8), GM clones (n = 13) and PG clones (n = 8) on day 12 of erythroid differentiation (mean ± s.d.). Multiple comparisons were assessed with one-way ANOVA with Scheffe's MCT. $**P < 0.01$, $****P < 0.0001$. **c** Flow cytometry analysis of the erythroid maturation markers CD71 and CD235 in the cells from (**a**).
